# The chemistry and morphology of *Vepris wadigo* (Rutaceae) a new Endangered tree of Kenyan coastal forest with the description of wadigin, a newly named alkaloid

**DOI:** 10.1101/2025.04.04.647269

**Authors:** Martin Cheek, W.R. Quentin Luke, Yanisa Olaranont, Moses Langat

## Abstract

Here we describe *Vepris wadigo* based on new collections and observations of the informally published *Diphasia* sp. A of the Flora of Tropical East Africa. *Vepris wadigo* is known with certainty only from Kaya Kinondo and Kaya Timbwa forests in Kwale District of coastal Kenya where it is known as Mchikoma (Digo language). A sterile specimen from the Usambara Mts of Tanzania suggests that the species may also occur there, but this needs verification with a fertile specimen.

*Vepris wadigo* is compared with the only other known diphasoid *Vepris* species of East Africa, *Vepris morogoroensis* of Tanzania. However, *Vepris wadigo* most closely resembles *Vepris stolzii* of Tanzania which differs in the 4-locular fruit.

The chemistry of *Vepris wadigo is* unique among the African species of *Vepris* which have been analysed, since acridones, quinolines and limonoids which characterise the genus were not detected. The six compounds which were characterised included compound **1** a benzamide newly described and named in this paper as wadigin (*N*-(2,6-dihydroxybenzoyl)-*O*-methyltyramine) which is an *O*-glucosylated derivative of riparin III, and four lignans and neolignans not previously found or very unusual in *Vepris*, namely heterotropan (**2**), asaraldehyde (**3**) a degradation product of compound **4**, *E-*asarone (**4)** and its isomer, *Z-* asarone (**5)**. Finally, Lupeol (compound **6**) which has previously been found in African *Vepris*.

*Vepris wadigo* is here provisionally assessed as Endangered, EN B1ab(i-iii)+B2ab(i-iii) since only two threat-based locations are known, with an area of occupation of 8 km^2^ using the IUCN preferred 4 km^2^ cell size, although the actual total area of the habitat occupied is 22.57 Ha as calculated with Google Earth Pro. Since there are only two points the extent of occurrence cannot be calculated. Satellite imagery has shown that the area of the habitat has slowly decreased in recent decades.

## Introduction

*Vepris wadigo* a new diphasioid species is described from two small coastal Kaya forests of Kenya and the phytochemistry of the leaves is reported. The species is based on an informally named taxon (material incomplete) published in the Flora of Tropical East Africa as *Diphasia* sp. A (Kokwaro 1982). This is supplemented by new observations and material recorded by the second author (see Fig. 1 & 2). The research for this paper was supported by preparation for a taxonomic revision of African *Vepris* Comm. ex A. Juss. by the first author, floristic work for conservation prioritisation in the surviving forests of Kenya and Tanzania by the second author, and study of the chemistry of *Vepris* species by the third and fourth authors.

**Fig. 1.**
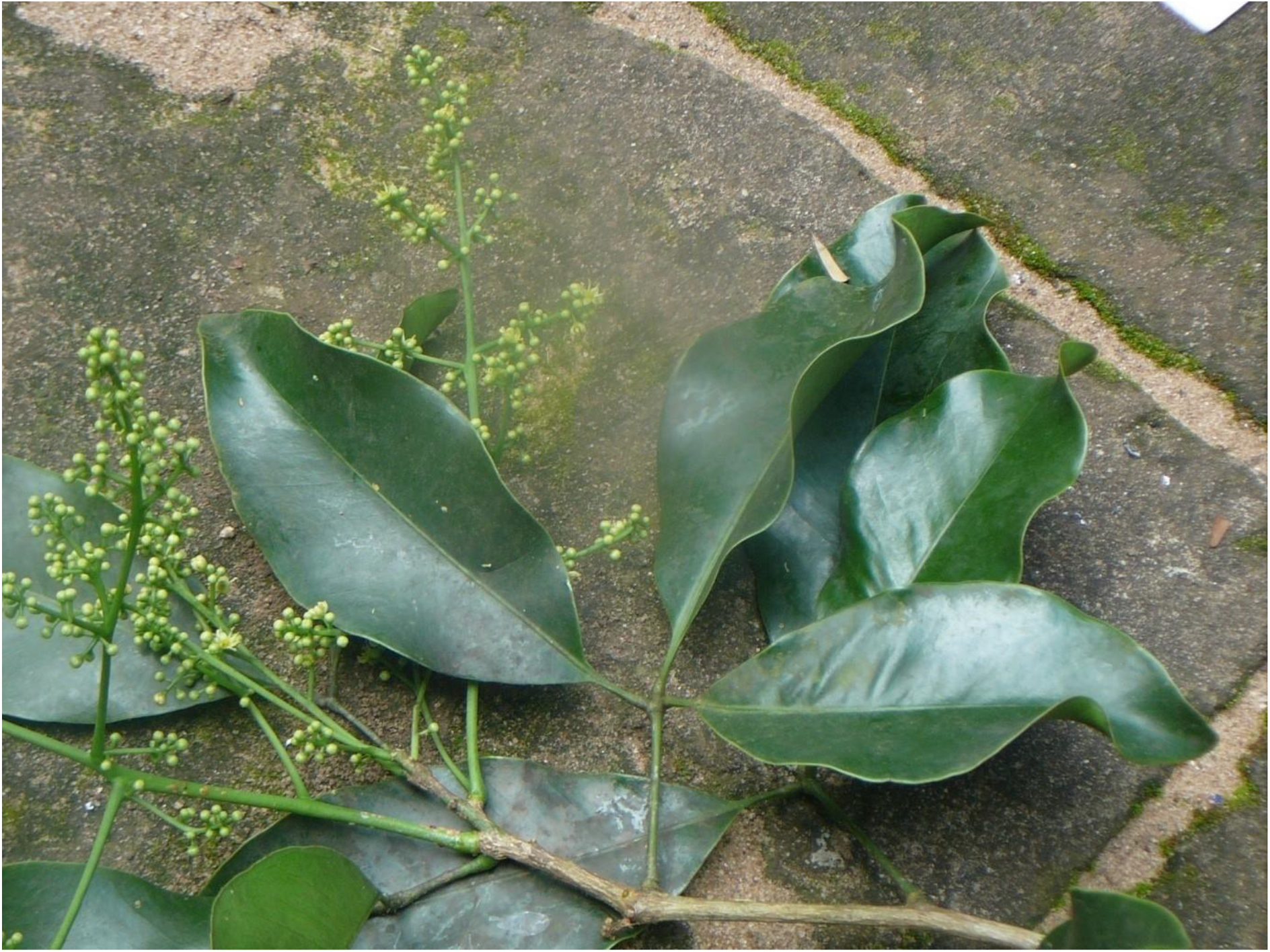
Vepris wadigo. Male flowering stem from Kaya Kinondo in August 2010. Photo by Elias Kimaru

**Fig. 2.**
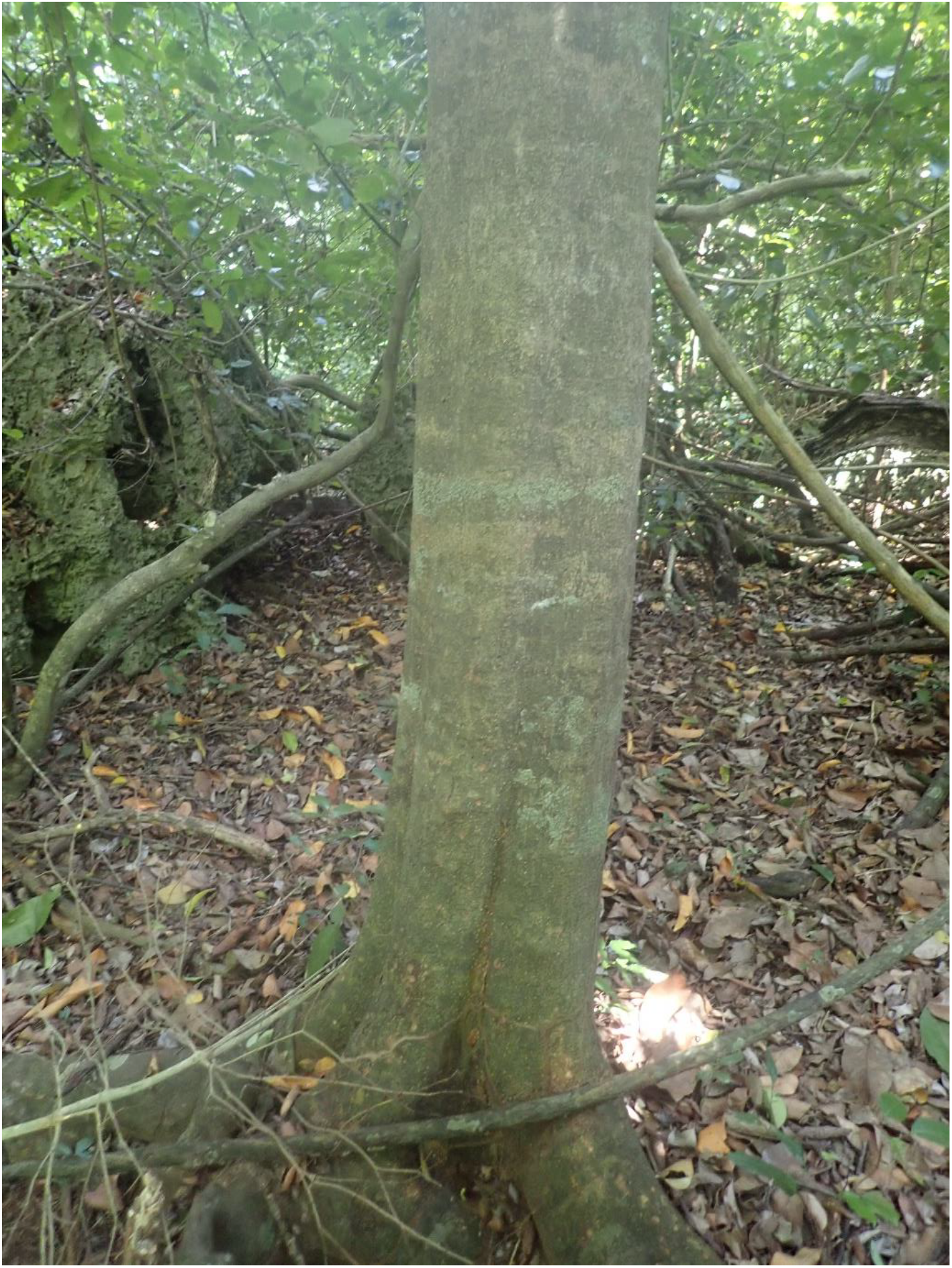
Vepris wadigo. Photo showing the trunk of a tree at Kaya Kinondo. Photo by Q. Luke August 2022.

*Vepris* (Rutaceae-Toddalieae), is a genus with 98 accepted species, 23 in Madagascar and the Comores and 74 in Continental Africa with one species extending to Arabia and another endemic to India (Plants of the World Online, continuously updated). The genus was last revised for tropical Africa by Verdoorn (1926). Founded on the Flore du Cameroun account of Letouzey (1963), nine new species were recently described from Cameroon (Onana & Chevillotte 2015; Cheek *et al*. 2018; Onana *et al*. 2019; Cheek & Onana 2021; Cheek *et al*. 2022a), taking the total in Cameroon to 24 species, the highest number for any country globally, followed by Tanzania with 19 species after recent additions by Cheek & Luke (2023) and by Ciambrone *et al*. (2024).

In continental Africa, *Vepris* are easily recognised. They differ from all other Rutaceae because they have digitately (1 –)3(– 5)-foliolate (not pinnate) leaves, and unarmed (not spiny) stems. The genus consists of evergreen shrubs and trees, predominantly of tropical lowland evergreen forest, but with some species extending into submontane forests and some into drier forests and woodland. *Vepris* species are often indicators of good quality, relatively undisturbed evergreen forest since they are not pioneers (Cheek & Luke 2023).

Species of *Vepris* in Africa extend from South Africa, e.g. *Vepris natalensis* (Sond.) Mziray, to the Guinean woodland in the fringes of the Sahara Desert (*Vepris heterophylla* (Engl.) Letouzey). Mziray (1992) subsumed the genera *Araliopsis* Engl., *Diphasia* Pierre, *Diphasiopsis* Mendonça, *Oricia* Pierre, *Oriciopsis* Engl., *Teclea* Delile, and *Toddaliopsis* Engl. into *Vepris*, although several species were only formally transferred subsequently (e.g. Harris 2000; Gereau 2001; Cheek *et al*. 2009; Onana & Chevillotte 2015). Mziray’s conclusions were largely confirmed by the molecular phylogenetic studies of Morton (2017) but Morton’s sampling was limited, identifications appeared problematic (several species appear simultaneously in different parts of the phylogenetic trees) and more molecular work would be desirable. Morton studied about 14 taxa of *Vepris*, all from eastern Africa. More recently Appelhans & Wen (2020) focussing on Rutaceae of Madagascar have found that the genus *Ivodea* Capuron is sister to *Vepris* and that a Malagasy *Vepris* is sister to those of Africa. However, most of the African species including all those of West and Congolian Africa, remain unsampled leaving the possibility open for changes to the topology of the phylogenetic tree when this is addressed.

Characteristics of some of the formerly recognised genera are useful today in grouping species. The “araliopsoid” species have hard, non-fleshy, subglobose, 4-locular fruit with 4 external grooves; the “oriciopsoid” soft, fleshy 4-locular syncarpous fruit; “oricioid” species are 4-locular and apocarpous in fruit; the fruits of “diphasioid” species are laterally compressed in one plane, bilocular and often bilobed at the apex; while “tecleoid” species are unilocular in fruit and 1-seeded, lacking external lobes or grooves. There is limited support for these groupings in Morton’s study.

Due to the essential oils distributed in their leaves, and the alkaloids and terpenoids distributed in their roots, bark and leaves, several species of *Vepris* have traditional medicinal value (e.g. Burkill 1997). Research into the characterisation and anti-microbial and anti-malarial applications of alkaloid and limonoid compounds in *Vepris* is active and ongoing (e.g. Atangana *et al*. 2017), although sometimes published under generic names no longer in current use, e.g. Wansi *et al*. (2008). Applications include as synergists for insecticides (Langat 2011). Cheplogoi *et al*. (2008) and Imbenzi *et al*. (2014) respectively list 14 and 15 species of *Vepris* that have been studied for such compounds. A review of ethnomedicinal uses, phytochemistry, and pharmacology of the genus *Vepris* was recently published by Ombito *et al*. (2020), listing 213 different secondary compounds, mainly alkaloids and furo- and pyroquinolines, isolated from 32 species of the genus, although the identification of several of the species listed needs checking. However, few of these compounds have been screened for any of their potential applications. Subsequently additional discoveries of compounds in *Vepris* have been published (Ciambrone *et al*. 2024; Langat *et al*. 2022; Mucaleque *et al*. 2024; Olaranont *et al*. 2023).

Recently, Langat *et al*. (2021) have published three new acridones and reported multi-layered synergistic anti-microbial activity from *Vepris gossweileri* (I.Verd.)Mziray, recently renamed as *Vepris africana* (Hook.f ex Benth.) Lachenaud & Onana (Lachenaud & Onana 2021).

There is no doubt that new compounds will continue to be discovered as chemical investigation of *Vepris* species continues.

## Materials and Methods - Morphology

The taxonomic study is based on herbarium specimens predominantly at EA, BM and K, by the first author and field observations of live material in Kenya by the second author. All specimens cited have been seen unless otherwise stated. The specimens were mainly collected using the patrol method e.g. Cheek & Cable (1997). Herbarium citations follow Index Herbariorum (Thiers *et al*. continuously updated), nomenclature follows Turland *et al*. (2018) and binomial authorities follow IPNI (continuously updated). Material of the new species was compared morphologically with material of all other African *Vepris*, principally at K, but also using material and images from BR, FHO, G, GC, HNG, P and YA. Herbarium material was examined with a Leica Wild M8 dissecting binocular microscope fitted with an eyepiece graticule measuring in units of 0.025 mm at maximum magnification. The drawing was made with the same equipment using a Leica 308700 camera lucida attachment. The description was made following the format of Cheek *et al*. (2022a) using terms from Beentje & Cheek (2003). Specimen location data is given as on the label of the specimens, understanding that the units formerly termed “Districts” in Kenya are currently termed Counties. For the extinction risk assessment, points were georeferenced using locality information from herbarium specimens. The conservation assessment was made using the categories and criteria of IUCN (2012). EOO was calculated with GeoCat (Bachman *et al*. 2011).

## Materials and Methods – Chemistry

The leaves were freeze-dried and ground to fine powder using a blender. The dried leaves were successively extracted using methylene chloride and methanol solvents. The methylene chloride extract was subjected to gravity column chromatography packed with 100% methylene chloride blend of silica gel Merck 9385 and eluted isocratically using 10% ethyl acetate in methylene chloride, collecting 35 mL. On the other hand, the methanol extract was subjected to gravity column chromatography packed with silica gel Merck 9385 suspended in 100% methylene chloride, and eluted isocratically using 5% methanol in methylene chloride, collecting 35 mL. The fractions were monitored using TLC and fractions with the same retention times were pooled.

Spectroscopic and spectrometric analysis were conducted as follows: 1D and 2D NMR spectra were recorded in CDCl_3_ and CD_3_OD on a 400 MHz Bruker AVANCE NMR instrument at room temperature. Chemical shifts (δ) are expressed in ppm and were referenced against the solvent resonances at δ_H_ 7.26 and δ_C_ 77.23 ppm for ^1^H and ^13^C NMR for CDCl_3_, at δ_H_ 4.87 and δ_C_ 49.15 ppm for ^1^H and ^13^C NMR for CD_3_OD. Accurate masses were recorded on a Thermo Scientific Orbitrap Fusion spectrometer. Purity of compounds was monitored *via* thin layer chromatography (TLC) using pre-coated aluminium-backed plates (silica gel 60 F_254_, Merck) and compounds were visualised by UV radiation at 254 nm and then using an anisaldehyde spray reagent (1% *p*-anisaldehyde:2% H_2_SO_4_: 97% cold MeOH) followed by heating. Final purifications used preparative thin layer chromatography (Merck 818133) and gravity column chromatography that was carried out using a 2 cm diameter column, which were packed with silica gel (Merck Art. 9385) in selected solvent systems.

## Results-Morphology

### Vepris wadigo

Cheek & Q. Luke sp. nov. Type: Kenya, Kwale District, Kaya Kinondo (Ngalani Kaya), fr. 16 July 1987, *Robertson & W*.*R*.*Q. Luke* 4867 (holotype K000875152, isotype EA). Figs. 1 – 3.

**Fig. 3.**
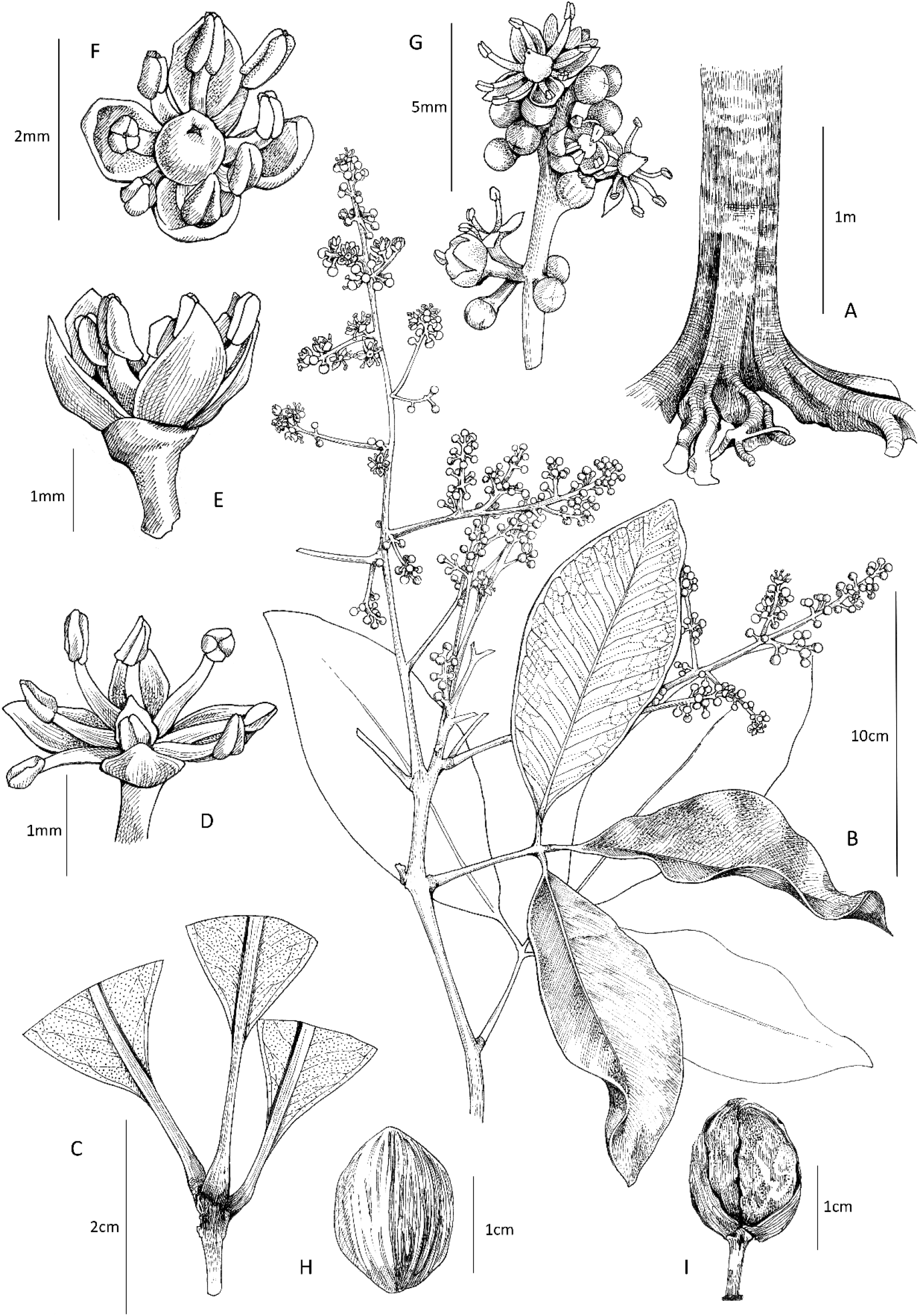
Vepris wadigo. **A**. habit, tree trunk; **B**. habit, flowering stem; **C**. part of leaf showing long petiolules; **D**. male flower, with pistillode; **E**. male flower, side view; **F**. female flower, with pistil; stem node and immature infructescence with leaf, showing winged petiole; **G**. portion of inflorescence; **H**. endocarp with seed, dorsal surface showing longitudinal fibre ridges; **I** side view of fruit with longitudinal groove dividing the two lobes. **A** from photo in Kaya Kinondo 2022 by WRQL (see Fig. 2); **B, D-E** from photo by Elias Kimaru in Kaya Kinondo 2010 (specimen mislaid, see Fig. 1); **C, H & I** from *Roberston & Luke* 4867 (K). Drawn by Meg Griffiths.

*Diphasia* sp. A Kokwaro (1982:32)

<u>Evergreen tree 4–25 m tall,</u> bark smooth, grey or pinkish grey. Leafy stems terete, drying pale brown, with 4–6 longitudinal ridges, 4–5.5 mm diam. at lowest leafy node, internodes 1.4 – 2.9(– 4) cm long, glabrous, lenticels sparse, concolorous, inconspicuous, (0.2 –)0.28 – 1.13 × 0.13 – 0.53 mm. <u>Leaves</u> alternate, spirally arranged, trifoliolate, long-petiolulate. Leaflets equal or subequal, the median sometimes slightly longer or shorter than the laterals, glossy dark green in life, drying brown-green, coriaceous, elliptic, or rarely, slightly obovate-elliptic, (12 –)15.4 – 22.7 x (4 –)5.6 – 7.2(– 8.4) cm, acumen short, apex rounded, 0 – 0.9 × 0 – 0.7 cm, less usually acute; base acute, slight asymmetric in lateral leaflets and decurrent as slender wings down the petiolules; secondary nerves (9 –)11 – 14 on each side of the midrib, arising at c. 45° from the midrib, rapidly curving outwards to c. 70° and then continuing straight towards the margin, via a looping inframarginal vein (c. 2mm from the margin) to the nerve above, tertiary nerves 1 – 3 between and parallel to the secondary nerves as intersecondaries, quaternary nerves inconspicuous; gland dots conspicuous in transmitted light, colourless, large, 0.08 – 0.15 mm dia., (5 –)7 – 9(– 15) per mm^2^ otherwise inconspicuous (not visible with naked eye) on surfaces, adaxial surface minutely pitted (microscope), concolorous; glands on abaxial surface grey; glabrous. Petiolules plano-convex, minutely winged, 1 – 2.5(– 3.7) cm long, 1 – 1.2(– 1.3) mm wide, basal pulvini 3 – 4 × 2(– 4) mm. Petioles terete, (4.3 –)6.2 – 11.2 cm long, 1.2 – 1.8 mm diam., glabrous. <u>Male Inflorescences</u> terminal and axillary in the distal 3 nodes, paniculate, 21 – 30 cm long, 8 – 12 cm wide, the partial-inflorescences themselves branched, glabrous; bracts inconspicuous, flowers single or more usually in fascicles or glomerules of 2 – 4-flowers, separated by internodes of 4 – 5 mm. <u>Pedicels</u> 1.1 – 2 mm long. Calyx saucer like, c. 0.5 – 0.7 mm wide, shallowly 4-lobed. Male flowers c. 2.5 mm wide at anthesis. Petals 4, white, ovate to oblong, c. 1.25 mm long, patent at anthesis. Anthers 8, subequal, outer whorl (alternate with petals) slightly shorter, inner (opposite petals), slightly longer c. 1.25 mm long. Pistillode conical, c. 0.7 mm diam., glabrous. <u>Female inflorescence</u> 6 – 12 cm long (Kokwaro 1982). Female flowers similar to male with larger pistils and slightly smaller anthers.

<u>Fruit</u> drupaceous, green, broadly ovoid-oblong, bilocular, 12 – 16 × 8 – 12 mm, 2-lobed laterally, barely laterally flattened, 2-seeded, surface glandular, glands pale yellow, orbicular c. 0.5 mm diam., pericarp leathery, mesocarp juicy; apex flattened to truncate, style base 2 – 3 × 0.3 mm. Fruiting pedicel 8 – 9 × 1 – 1.5 mm. <u>Seed</u> plano-convex, elliptic in outline, 12 – 13 × 9 × 5.5 mm, encased in a golden-yellow fibrous leathery endocarp, the dorsal surface with strongly raised longitudinal ridges.

### RECOGNITION

Differing from *Vepris morogoroensis* (Kokwaro) Mziray of Tanzania, the only other bilobed and bi-locular fruited (diphasoid) species published from E Africa, in being glabrous, and having terete petioles (vs minutely pubescent, petioles canaliculate), in the long-petiolulate leaflets (vs sessile or subsessile), and in the inflorescences as long or longer than the leaves, male flowers with 8 stamens (vs inflorescences shorter than the petioles, male flowers with 4 stamens).

### DISTRIBUTION & HABITAT

Kenya, Kwale County, Kinondo & Timbwa Kaya forests (possibly also Usambara Mts, see Notes). Evergreen or semi-deciduous forest on Coral Rag with *Milicia, Antiaris, Lannea welwitschii, Terminalia catappa, Diospyros ferrea, Calophyllum, Sorindeia* and *Drypetes*.

### SPECIMENS EXAMINED. KENYA

Kwale District, Kinondo (Ngalani Kaya) Forest, K7, 4° 23’ 30” S 39° 33’ 30” E, 5 m, fr., 15 Feb 1977, *R*.*B. Faden & A*.*J. Faden* 77/389 (K000875151!); ibid, Kaya Kinondo, K7, 04.235 S 39.325 E, 0 – 10 m, fr., 16 Jul 1987, *S*.*A. Robertson & W*.*R*.*Q. Luke* 4867 (holo K000875152!; iso EA); ibid, Kaya “Mgawani” (Timbwa) 7 m, st., 4 Jan 1992, *W*.*R*.*Q. Luke* 3035 (EA, K000875150!).

### CONSERVATION

*Vepris wadigo* appears endemic to Kaya Kinondo and the nearly adjacent Kaya Timbwa. “*Kaya Kinondo is a 30-hectare forest located in the Diani Beach area south of Mombasa. The local Digo community has formed the Kaya Kinondo Ecotourism project that offers controlled access into this pristine and untouched forest. The forest is still used for rituals and sacrificial offerings by the local Digo tribe. A walk through the forest, which forms a rich ecosystem with 187 species of plants, some of which are rare and a few, endemic to the area. There are also animals, mostly primates (Angolan Colobus, Sykes, Vervet and Bushbabies), dik-diks, and Suni antelope. You will also have the opportunity to visit Kinondo Village and interact with members of the local community including the medicine man. The project was funded in 2001 with the initial support from WWF and NMK and the community has seen better protection of the Kaya which has realized socio-economic benefits*” (https://www.musement.com/uk/mombasa/kaya-kinondo-sacred-forest-half-day-tour-from-mombasa-south-coast accessed 23 July 2022)

According to the notes on *R & A Faden* 77/789 (K), the species was frequent at Kaya Kinondo in 1977. Kaya Kinondo sacred forest is now known to contain 231 species of plant (WRQ Luke personal records 2022). The forest currently measures 550 m north to south, and the maximum east to west is 420 m, the minimum 230 m, giving a polygon measured as 16 Ha (Luke pers. obs. 2022). The forest is only 10 m distant from the shore of the Indian ocean (Maxar Technologies image dated 17 Jan. 2017, viewed 23 July 2022 on Google Earth Pro). Using the time slider on Google Earth Pro shows that since October 2001 the forest has been reduced in extent in the north and south by up to 140 m in length, and the former northernmost part of the forest is now walled, as though for a building plot. New ecotourism buildings have been built inside the new northern boundary of the forest. Holiday residences with pools have been built adjoining the forest to the west and east since 2001. While the perimeter of the current forest boundary seems well-defined and intact and the canopy of the major portion is also mainly intact, some small open areas and fallen trunks can be seen within, but these are due to natural windthrow from storms (WRQL pers. obs. 2022).

At Kaya Timbwa (erroneously sometimes as referred to as Mgawani), the second locality, Luke (*Luke* 3835, K) considered the forest to be possibly secondary, post-cultivation, citing presence of large *Terminalia catappa*. Kaya Timbwa lies about 650 m SSW of Kaya Kinondo, extending c. 490 m approximately east to west, and about 200 m north to south. Over the last 20 years its area has been stable but some large trees have been lost from the eastern edge, and the numbers of buildings at its northern edge has increased. We infer that these two forests were once continuous but became fragmented by clearance of the intervening forest.

We assess *Vepris wadigo* as Endangered, EN B1ab(i-iii)+B2ab(i-iii) since only two threat-based locations are known, with an area of occupation of 8 km^2^ using the IUCN preferred 4 km^2^ cell size, although the actual total area of the habitat occupied is 22.57 Ha as calculated with Google Earth Pro. Since there are only two points, extent of occurrence cannot be calculated but is estimated as about the same as the AOO. The main threat appears to be continued reduction of the area of surviving habitat over time, since although this has been slow and small, EOO, AOO and extent of habitat have been reduced. If extreme fragmentation is accepted, the species could instead be assessed as Critically Endangered.

### NOTES

*Vepris wadigo* has been categorized previously as a *Diphasia* due to its stated two-lobed fruit (Kokwaro 1982 as *Diphasia sp*. A). In terms of its fruit, it is undoubtedly most similar to *Vepris morogoroensis*. Vegetatively and in stamen number, the two differ greatly (see the diagnosis) and cannot be confused. In almost all other respects *Vepris wadigo* more closely resembles and is likely to be confused with *Vepris stolzii* I. Verd. Both species are glabrous, have coriaceous long-petiolulate leaflets, large terminal panicles of 8-stamened flowers. They differ in that the last species has 4-locular, tetragonal fruit (vs 2-locular, laterally flattened fruit). Further herbarium collections of *Vepris wadigo* are desirable. While flowers are known (Fig. 1, description) the specimen concerned was mislaid.

### ETYMOLOGY

The specific epithet wadigo means “of the Digo”. The Digo are one of the nine tribes of the Mjikenda of coastal Kenya, who maintain their ancestral Kaya forests for traditional purposes. The Kaya forests are also globally important for conservation of plant species, with many species restricted to them. *Vepris wadigo* is only known with complete certainty from two Kaya forests of the Digo in Kwale County, Kenya (but see notes below). **VERNACULAR NAMES & USES**. Mchikoma (Digo). Used to chase out evil spirits. The patient is steamed under a blanket with the leaves of this species (Elias Kimaru pers. comm. to W.R.Q. Luke in 2022).

### NOTES

The species was first described by Kokwaro (1982) as *Diphasia* sp. A, on the basis of the single specimen *R &A Faden* 77/389.

*Luke & Muir* 6084 (K!) from Mbomole Hill (5.10 S, 38.63 E), E. Usambaras, Tanzania, 11 Dec 1999, 900 m alt. closely resembles *Vepris wadigo*, differing in the slightly larger (9 cm wide), obovate leaflets, which lack the gland pits on the adaxial surface (the glands instead are raised), which are thinly papery (vs coriaceous), and which have conspicuous, reticulate quaternary nerves (vs inconspicuous). However, these differences might be explained by the specimen being from a 2 m tall sapling, not the crown of a mature tree as are the Kenyan specimens of *Vepris wadigo*. However, an element of doubt remains, although the authors judge that it is 90% certain that this specimen represents a third location for *Vepris wadigo*. It is just possible, although unlikely, that this represents a similar, but different species. Confirmation of the identification is desirable by collection of a fertile specimen or one with leaves from the crown at the Usambara location before we can be certain of this extension of the range.

## Results-Chemistry

The leaves of *V. wadigo* gave six compounds (**1**-**6** see **1**-**5** in Fig. 4 above). Compound **1**, wadigin, is an undescribed *O*-glucosylated derivative of riparin III (*N*-(2,6-dihydroxybenzoyl)-*O*-methyltyramine) previously reported from unripe fruits of a known medicinal plant, *Aniba riparia* (Nees)Mez (Lauraceae, northern S. America, Barbosa-Filho *et al*. 1987). Compound **2**, heterotropan, is a known neolignan previously isolated from *Asarum fauriei* var. *takaoi* (F.Maek.) T.Sugaw (formerly *Heterotropa takaoi* (F.Maek.)F.Maek. Aristolochiaceae, Japan, Yamamura *et al*. 1978) and *Piper cubeba* L.f. (Piperaceae, Myanmar to Indonesia, Badheka *et al*. 1987). Compound **3**, asaraldehyde is a degradation product of compound **4**, *E-*asarone, and its *Z-*isomer, compound **5**, *Z-*asarone, also isolated from the *Asarum sp*. Compound **2** is a 2+2, cyclo-addition product of **4** or **5**. Lupeol was also isolated from this plant as compound **6**.

**Fig. 4.**
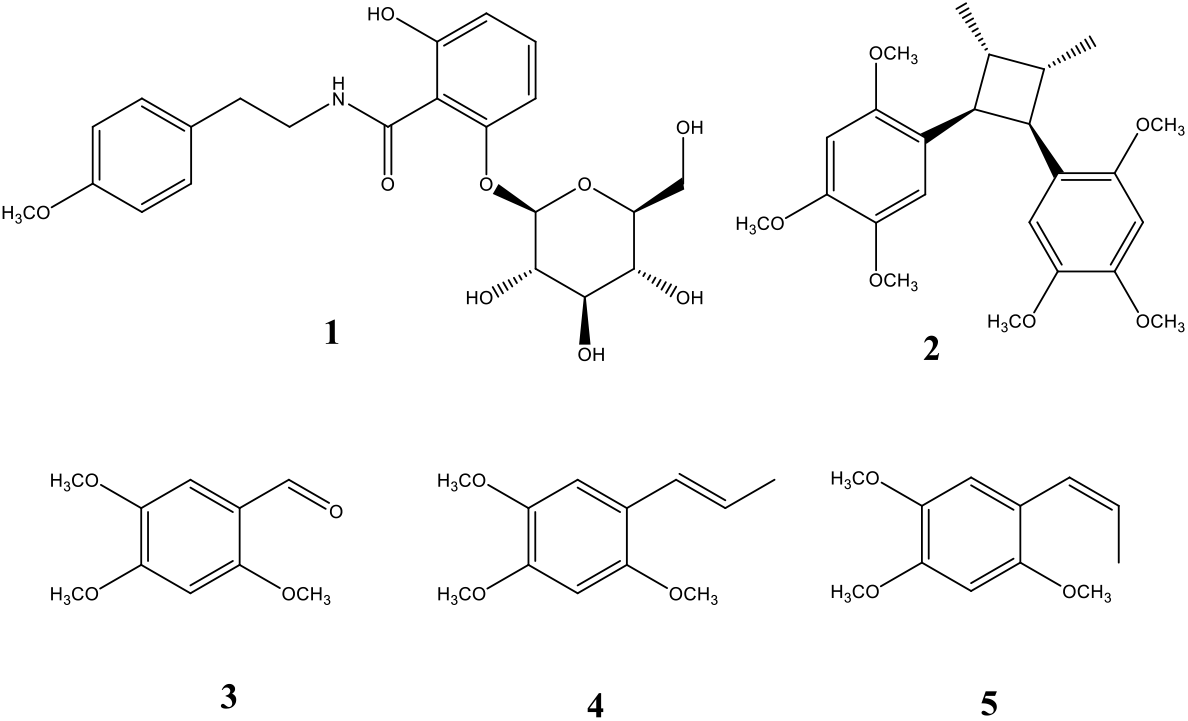
Structures **1-5** of compounds isolated from *Vepris wadigo* including a new *O*-glucosylated derivative of riparin III, herein, named wadigin (structure **1**).

The ^1^H NMR spectrum for compound **1** showed five proton resonances in the aromatic region with two characteristic splitting systems, also supported by COSY spectrum. The splitting systems were AA’XX’ and ABX, for a para-substituted phenyl ethyl group, and disubstituted benzoic acid derivative. A pair of two proton resonances at δ_H_ 7.20 (d, 9.0 HZ) and δ_H_ 6.87 (d, 9.0 HZ), the methoxy proton resonance at δ_H_ 3.79 (s), and a pair of two double doublets for methylene proton resonances at δ_H_ 2.86 (dd 7.0, 7.0 Hz) and 2.27 (dd 7.0, 7.0 Hz), and δ_H_ 3.70 (m) and δ_H_ 3.50 (m) for a substituted ethyl group. The de-shielded methylene proton resonances at δ_H_ 3.70 (m) and δ_H_ 3.50 (m) showed a correlation in the HMBC spectrum with a carbonyl carbon resonance at δ_C_ 177.5 attributable to benzoic acid ester. The ABX proton resonance was δ_H_ 7.28 (dd, 8.4, 9.0 Hz), δ_H_ 6.84 (dd, 9.0, 1.0 Hz) and δ_H_ 6.61 (dd, 8.4, 1.0 Hz). In addition, proton resonances for a glycosyl group were observed, and were assignable to a glucose group. The anomeric proton resonance was at δ_H_ 4.94 (d, 7.9 Hz), and the coupling constant supported a β-anomer (Table 1).

**Table 1:** ^13^C and ^1^H NMR chemical shifts for compound **1** (wadigin)

| No. | $^{13}\text{C}$ NMR | $^1\text{H}$ NMR |
| --- | --- | --- |
| 1 | 105.2 | - |
| 2 | 162.3 | - |
| 3 | 113.2 | 6.61 dd 8.4, 1.0 |
| 4 | 134.2 | 7.28 dd 8.4, 9.0 |
| 5 | 107.6 | 6.84 dd 9.0, 1.0 |
| 6 | 157.0 | - |
| 7 | 177.5 | - |
| 1' | 129.1 | - |
| 2'/6' | 130.6 | 7.20 d 9.0 |
| 3'/5' | 114.7 | 6.87 d 9.0 |
| 4' | 158.5 | - |
| 7' | 35.3 | 2.86 dd 7.0, 7.0 |
|  |  | 2.27 dd 7.0, 7.0 |
| 8' | 41.9 | 3.70 m |
|  |  | 3.50 m |
| NH | - | - |
| OCH<sub>3</sub> | 55.4 | 3.79, s |
| 1'' | 103.6 | 4.94 d 7.9 |
| 2'' | 71.1 | 3.32 m |
| 3'' | 77.9 | 3.44 m |
| 4'' | 73.6 | 3.36 m |
| 5'' | 78.5 | 3.46 m |
| 6'' | 62.3 | 3.91 m |
|  |  | 3.71 m |

## Discussion

Several other highly range restricted species occur only in Kaya Kinodo or one other forest, or are endemic to other Kaya forests nearby in coastal Kenya. Recently published examples include *Croton kinondoensis* G.W.Hu, Ngumbau & Q.F.Wang (Euphorbiaceae) which is only known globally from Kaya Kinondo (Ngumbau *et al*. 2020) and *Grevea eggelingii* Milne Redh.var. *keniensis* Verdc. (Montiniaceae, Verdcourt 1996) is near endemic to Kaya Kinondo (with a second location in Tanzania).

*Premna mwadimei* Ngumbau & G.W.Hu (Lamiaceae, Ngumbau *et al*. 2021) is endemic to the Kaya forest of Cha Simba, while *Lukea quentinii* Cheek & Gosline (Annonaceae, Cheek *et al*. 2022b) is endemic to Kaya Ribe, *Afrothismia baerae* Cheek (Afrothismiaceae, Cheek 2004a, Cheek *et al*. 2024), *Keetia lukei* Bridson (Rubiaceae, Bridson 1994) and a species of *Uvariodendron* (Dagallier *et al*. 2021) are all endemic to one to two other Kaya forests. The *Croton* is especially interesting as its affinities are with species of Madagascar rather than continental Africa (Ngumbau *et al*. 2020).

According to our analysis, *Vepris wadigo* in this phytochemical study, entirely lacks acridones, quinolines or limonoids, making it unusual, in fact so far, unique in the genus which chemically is otherwise characterised by the presence of these compounds (Ombito *et al*. 2020). In fact, apart from compound **6**, Lupeol, which is widespread in species of African *Vepris*, compounds **1-5** are previously unknown in *Vepris*, although derivatives of compound **1** (wadigin) are known in two other African species of the genus, and other lignans and neolignans (compounds 2-5) can also occur in African *Vepris* (Ombito *et al*. 2020). A possible explanation for the mismatch between the chemistry of *Vepris wadigo* and all other known continental African species of *Vepris* which have been investigated for chemistry, may be that as with *Croton kinondoensis* (see above), the affinities of this *Vepris* may be with the species of *Vepris* in Madagascar. Chemistry of the species in Madagascar is very little researched, and it is possible that when this is addressed, matches for *Vepris wadigo* will be found. Molecular phylogenetic studies, ideally nuclear phylogenomic with multiple genes are needed across the species of both Africa and Madagascar to decide on the affinities of this species. Studies of the chemistry of more species of *Vepris* from Madagascar would also enable this hypothesis to be tested.

*N*-[2-(3,4-Dimethoxyphenyl)ethyl]-2-(β-*D*-glucopyranosyloxy)-6-hydroxybenzamide, a methoxylated derivative of (**1**) has been prepared in the lab and determined to be an antidiabetic agent and SGLT2 inhibitor (Washburn *et al*. 2001). Riparin III, of which compound **1** (wadigin, (*N*-(2,6-dihydroxybenzoyl)-*O*-methyltyramine)) is an *O*-glucosylated derivative, is an interesting compound that has been demonstrated to have antioxidant, anti-inflammatory and antidepressant effects. The antibiofilm properties of Riparin II-type benzamides are considered to give them potential as new drugs targeting dermatophytes by inhibiting the Ssu1 protein (da Rocha *et al*. 2024). *N,N*-dimethyl 4-methoxyphenylethylamine, a derivative of compound **1** herein has been previously reported from *Vepris simplicifolia* (now *Vepris viridis* (I,Verd.) Cheek et al., see Cheek *et al*. 2025) by Badger *et al*. (1963). Candicine, a *N,N,N*-trimethyltyramine, also derived from tyrosine, was reported from *V. heterophylla* (Waterman *et al*. 1978), therefore our report of **1** in *Vepris wadigo* represents only the third report of this rare group of compounds in the genus *Vepris*. The presence of lignans in *Vepris* has been reported, including in *V. uguenensis*, by Cheplogoi *et al*. (2008) and *V. glomerata* by Kiplimo & Koorbanally (2012). Presence of phenyl propananoid, (*E*)-anethole was reported from *V. elliotii* I.Verd. by Poitou *et al*. (1995).

## Conclusions

The new species published in this paper is restricted to Kaya forests of coastal Kenya which are part of the Eastern Arc Mountains and Coastal Forests (EACF) of Tanzania and southern Kenya. The EACF form an archipelago-like phytogeographical unit well-known for high levels of species endemism in many groups of organisms (Gereau *et al*. 2016). Apart from the coastal forests that include *Vepris wadigo*, the better-known blocks are the Nguru Mts, the Udzungwa Mts, the Uluguru Mts, and the Usambara Mts. In herbaceous groups such as the Gesneriaceae, over 50% of the taxa (23 endemic species and a further nine endemic taxa) for East Africa (Uganda, Kenya and Tanzania) are endemic to the EACF (Darbyshire 2006) and in the Acanthaceae, there are numerous endemic species in multiple genera endemic to the EACF, e.g. *Stenandrium warneckei* (S.Moore) Vollesen, *Isoglossa bondwaensis* I. Darbysh., *Isoglossa asystasioides* I. Darbysh. & Ensermu and *Sclerochiton uluguruensis* Vollesen (Darbyshire 2009; Darbyshire *et al*. 2010; Darbyshire & Kelbessa 2007). In terms of documented plant species diversity per degree square, the EACF are second only in tropical Africa to Southwest Cameroon in the Cross-Sanaga Interval of West-Central Africa (Barthlott *et al*. 1996; Cheek *et al*. 2001).

The EACF include the sole representatives of plant groups otherwise restricted on the continent to the forests of Guineo-Congolian Africa, e.g. *Afrothismia* Schltr. *Ancistrocladus* Wall. and *Mischogyne* Exell (Afrothismiaceae, Cheek & Jannerup 2006; Ancistrocladaceae, Cheek *et al*. 2000, Cheek 2000, Léonard 1984; Annonaceae, Gosline *et al*. 2019). Several EACF forest genera have even more disjunct distributions, being found only in the Cross-Sanaga Interval and in the EACF and not in between, e.g. *Zenkerella* Taub. (Leguminosae) and *Kupea* Cheek (Triuridaceae Cheek 2004b). Extensive forest clearance within the last 100–150 years has removed forest from some areas entirely, and reduced forest extent greatly in others (this paper and Ndang’ang’a *et al*. 2007).

The provisional extinction risk assessment of *Vepris wadigo* of Endangered EN B1ab(i-iii)+B2ab(i-iii) is a cause for concern. Thankfully, there is a level of local community protection by the indigenous people of SE Kenya of their Kaya forests. However, the future for the species seems fragile.

Until species are scientifically named, it is difficult for an IUCN conservation assessment to be published (Cheek *et al*. 2020). Most new species to science published today, such as in this paper, are range-restricted, meaning that they are almost always automatically threatened, although there are exceptions, such as the widespread *Vepris occidentalis* Cheek & Onana (Cheek *et al*. 2019). Documented extinctions of plant species are increasing (Humphreys *et al*. 2019) and recent estimates suggest that as many as 2/5 of the world’s plant species are now threatened with extinction (Nic Lughadha *et al*. 2020) and 3/4 of those published today are already threatened (Brown *et al*. 2023). Global extinctions of African plant species continue apace e.g. in West Africa (*Inversodicraea pygmaea* G.Taylor and *Saxicolella deniseae* Cheek in Guinea, Cheek *et al*. 2017, 2022c), in Central Africa (e.g. Cheek *et al*. 2021) as well as in East Africa. In Tanzania, at the foot of the Udzungwa Mts, the achlorophyllous mycotrophs *Kihansia lovettii* Cheek and *Kupea jonii* Cheek (Triuridaceae, Cheek 2004b) are likely extinct as a result of the placement of the Kihansi hydroelectric dam, not having been seen since construction in 1994 (31 years ago), despite targeted searches. Although not directly threatened by development, another mycotroph, this time in another coastal forest fragment of SE Kenya, *Afrothismia baerae* Cheek (Afrothismiaceae, Cheek 2004a) has also not been found despite monitoring in the last 10 years. If future extinctions are to be avoided, improved conservation prioritisation exercises are needed such as Important Plant Area programmes (Darbyshire *et al*. 2017), supported by greater completion of Red Listing, although this can be slow and problematic (Bachman *et al*. 2019) and, globally, only 21 – 26 % of plant species have conservation assessments (Bachman *et al*. 2018). Where possible, as an insurance policy, seed banking and cultivation of threatened species in dedicated nurseries are urgent. Above all, completion of botanical taxonomic inventories is needed to feed into these exercises, otherwise we will continue to lose species before they are even discovered for science, and certainly before they can be investigated for their potential for beneficial applications. New compounds to science with high potential for humanity are being discovered in *Vepris* species each year (e.g. potent antimicrobial compounds in *Vepris africana*, Langat *et al*. 2021) and it is possible that wadigin, characterised and named in this paper, might also be found in future to have medicinal applications. Such discoveries will not be possible if species extinctions are allowed to continue.

## Supporting information

Supplementary information chemistry of Vepris wadigo

## Acknowledgements

WRQL would like to thank respectfully Chitiro Mwaukonde of Kaya Kinondo for supporting visits there, and discussions. Elias Kimaru is thanked for sharing information on the local name and use, and photos included in this paper.

We thank Nina Davies for facilitating the loans and working spaces needed for this paper in the Kew Herbarium. Greta Buinovaskaja and Shigeo Yasuda assisted in the early stages of this project collecting morphological data. Janis Shillito typed the manuscript.

The authors declare that they have no conflict of interest.

