## Supplementary information chemistry of Vepris wadigo for "The chemistry and morphology of *Vepris wadigo* (Rutaceae) a new Endangered tree of Kenyan coastal forest with the description of wadigin, a newly named alkaloid"

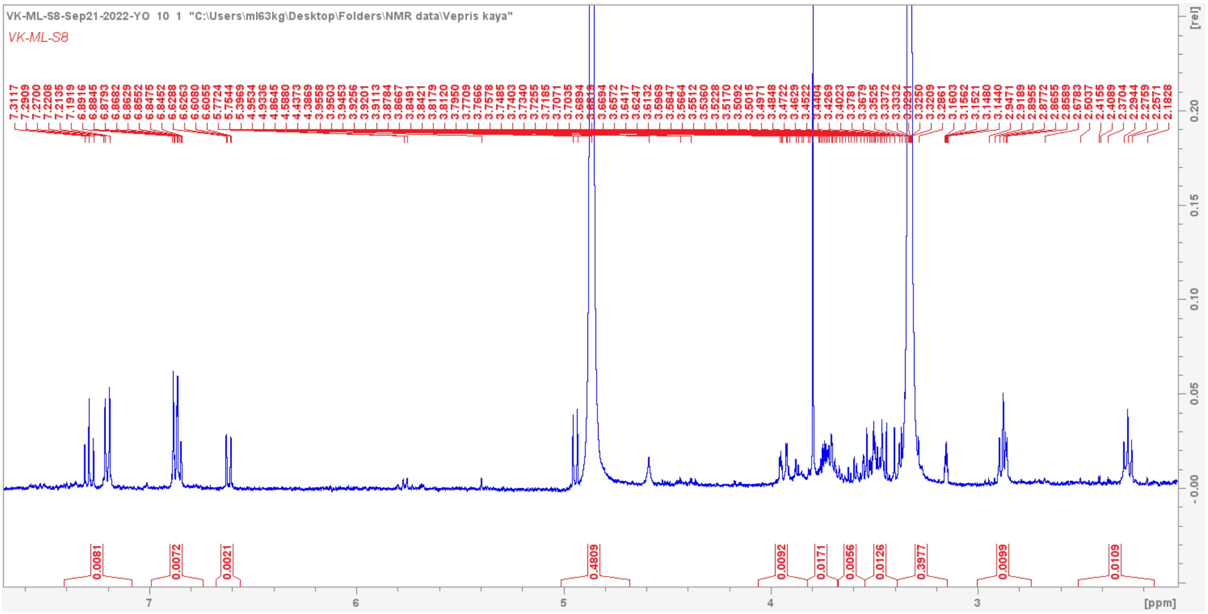

1H NMR for compound 1

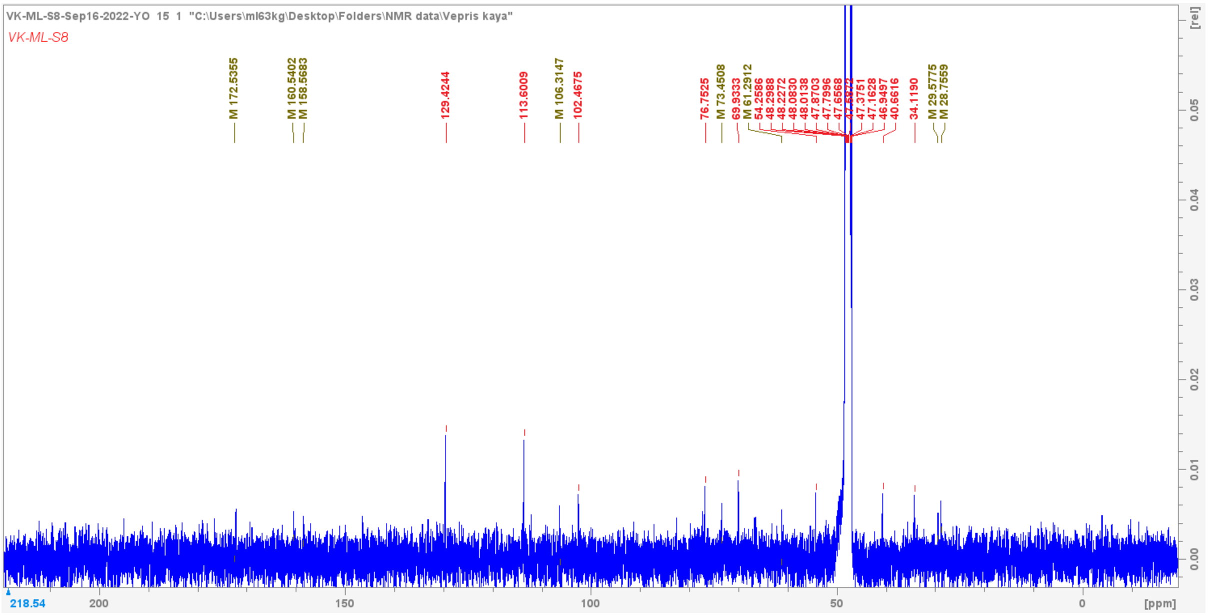

13C NMR for compound 1

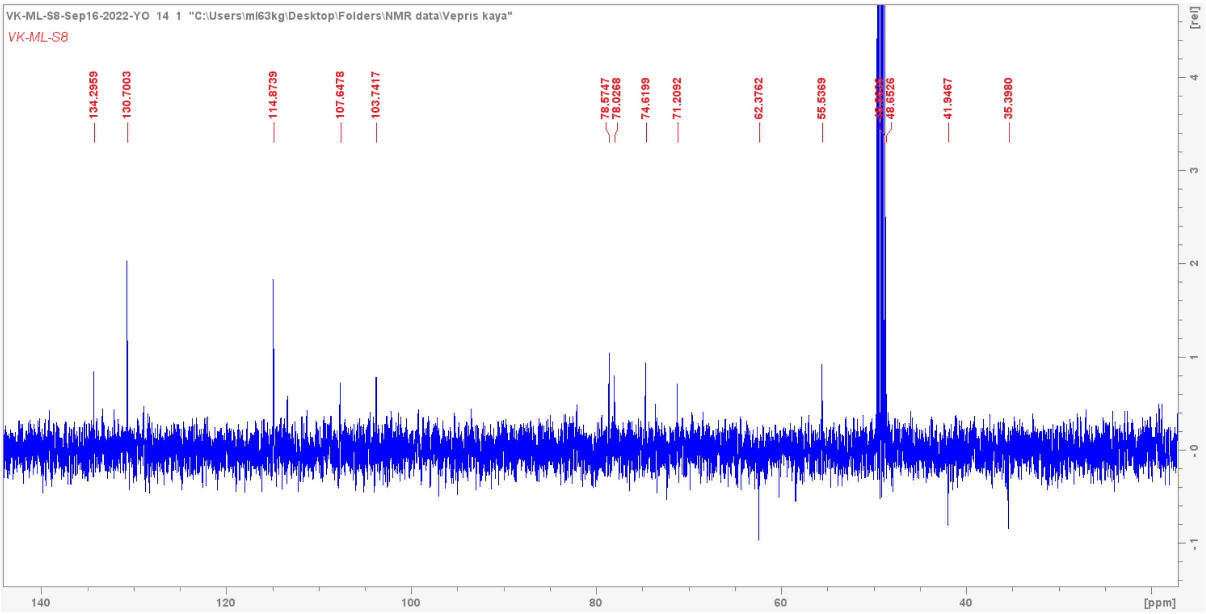


DEPT spectrum for compound 1

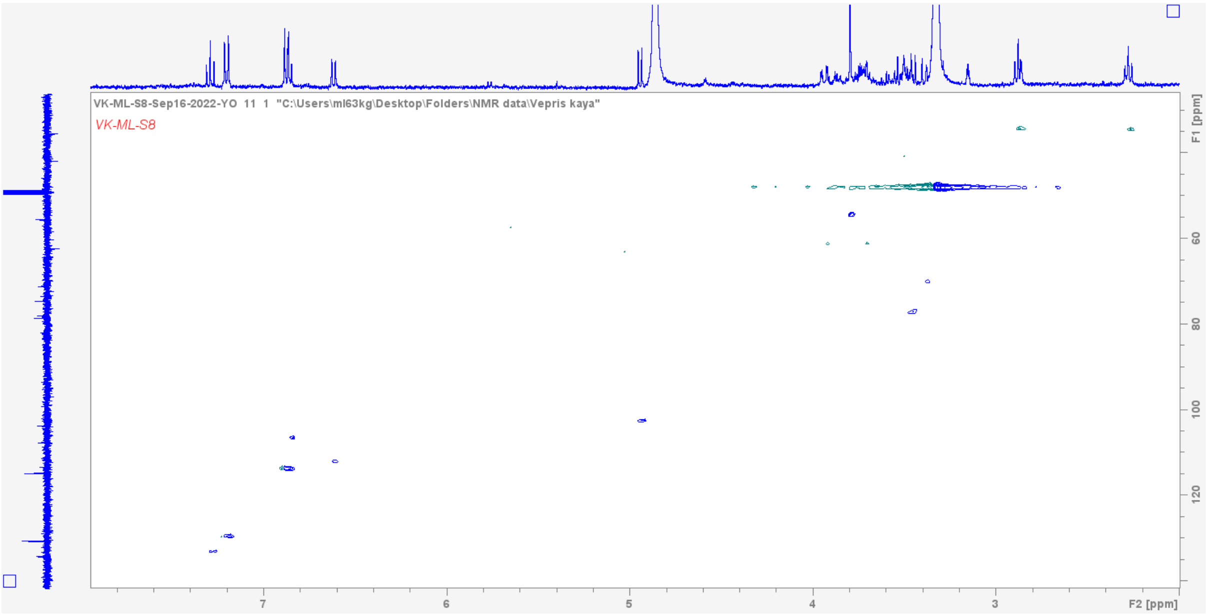


HSQCDEPT for compound 1

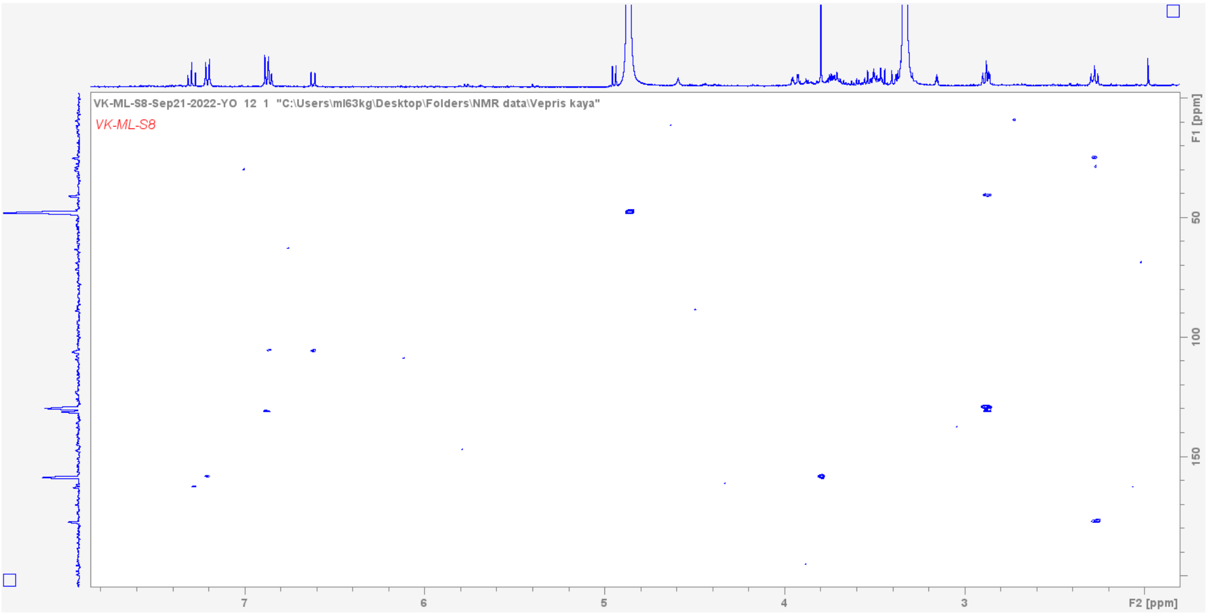


HMBC spectrum for compound 1

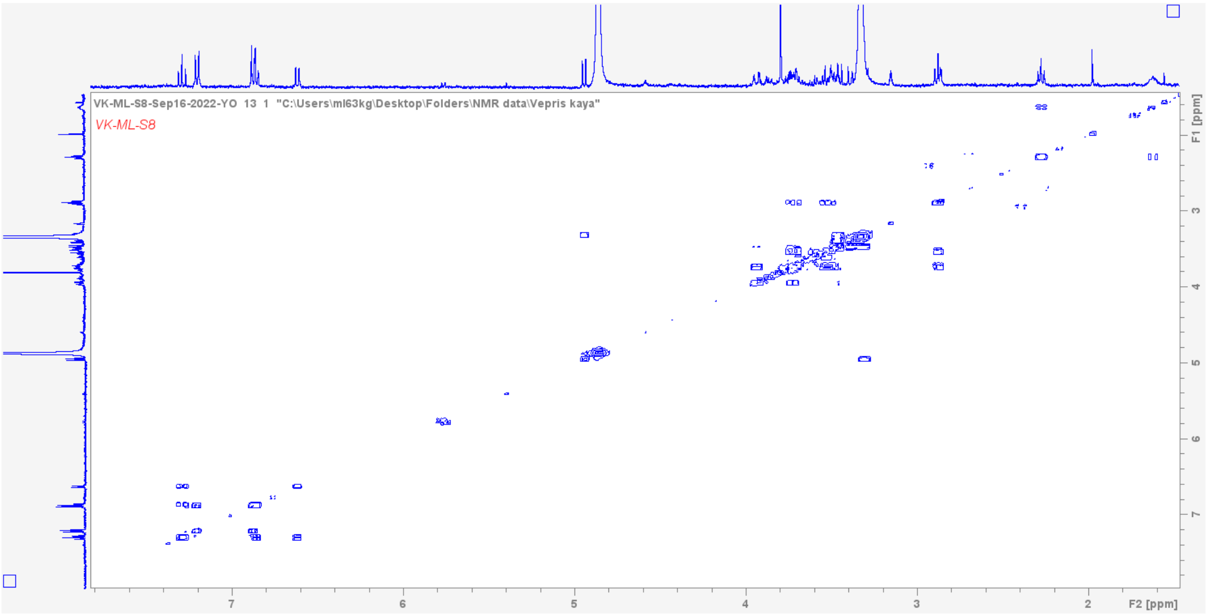


COSY spectrum for compound 1

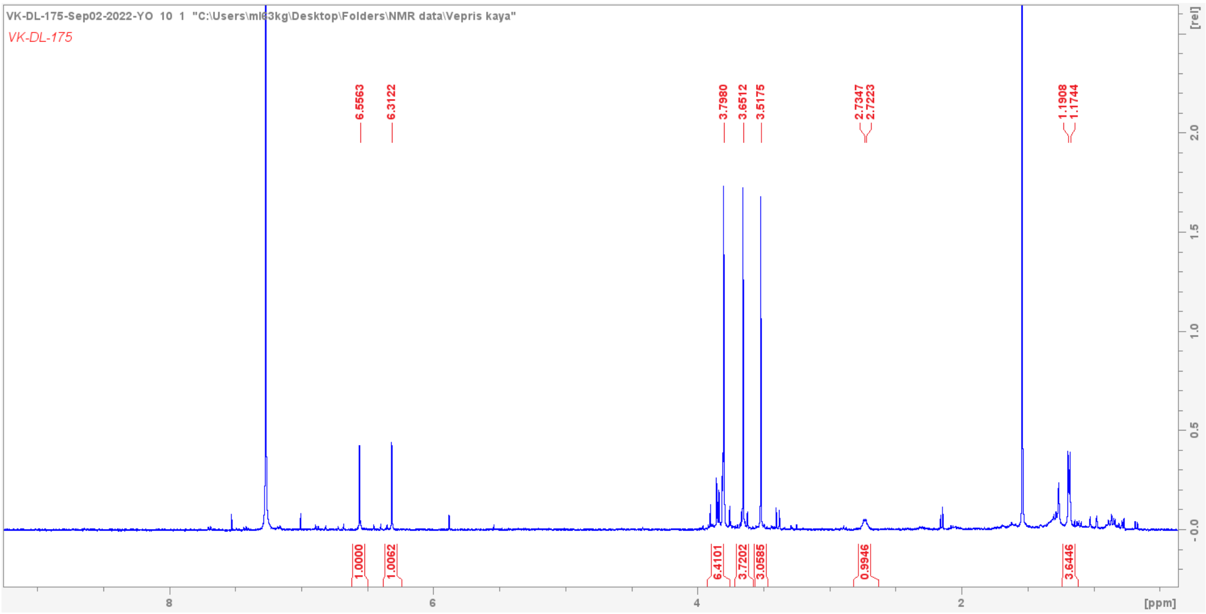


1H NMR for compound 2

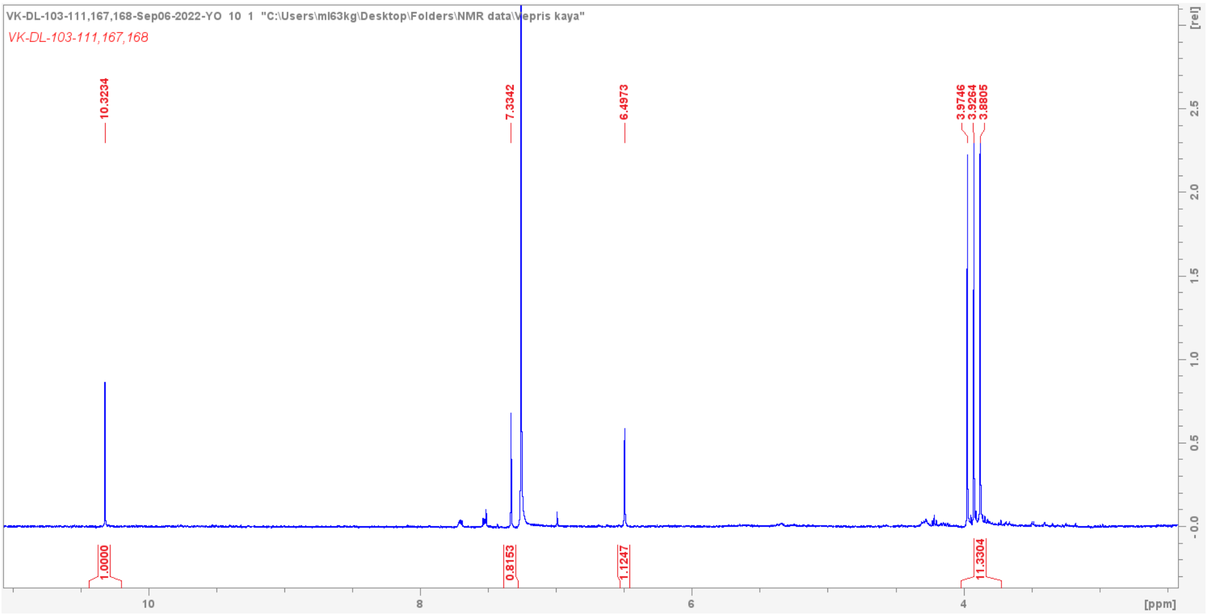


1H NMR for compound 3

Fraction 109 – 116 and
